# A Novel Framework for *In-vivo* Diffusion Tensor Distribution MRI of the Human Brain

**DOI:** 10.1101/2022.08.15.503969

**Authors:** Kulam Najmudeen Magdoom, Alexandru V. Avram, Joelle E. Sarlls, Gasbarra Dario, Peter J. Basser

## Abstract

Neural tissue microstructure plays an important role in developmental, physiological and pathophysiological processes. Diffusion tensor distribution (DTD) MRI helps probe heterogeneity at the mesoscopic length scale, orders of magnitude smaller than the nominal MRI voxel size, by describing water diffusion within a voxel using an ensemble of non-exchanging compartments characterized by a probability density function of diffusion tensors. In this study, we provide a new framework for acquiring tensor encoded diffusion weighted images (DWIs) and estimating DTD from them for *in-vivo* human brain imaging. We interfused pulsed field gradients (iPFG) in a single spin echo to generate arbitrary b-tensors of rank one, two, or three without introducing concomitant gradient artifacts. Employing well-defined gradient pulse duration and mixing/diffusion times in our diffusion preparation, we show that iPFG retains salient features of traditional multiple-PFG (mPFG) sequence while overcoming some of its implementation issues thereby extending its applications beyond DTD MRI. We assume DTD is a maximum entropy tensor-variate normal distribution whose tensor random variables are constrained to be positive definite (CNTVD) to ensure their physicality. In each voxel, the second-order mean and fourth-order covariance tensors of the DTD are estimated using a Monte Carlo method that synthesizes micro-diffusion tensors with corresponding size, shape and orientation distributions to best fit the measured DWIs. From these tensors we obtain the mean diffusivity (MD) spectrum, spectrum of diffusion tensor shapes, microscopic orientation distribution function (*µ*ODF), and microscopic fractional anisotropy (*µ*FA) which disentangle the underlying heterogeneity within a voxel. Using DTD derived *µ*ODF, we introduce a new method to perform fiber tractography capable of resolving complex fiber configurations. The results obtained in the live human brain showed microscopic anisotropy in various gray and white matter regions and skewed MD distribution in cerebellar gray matter not observed previously. DTD MRI tractography captured complex white matter fiber organization consistent with known anatomy. DTD MRI also resolved some degeneracies associated with diffusion tensor imaging (DTI) and identified the source of microscopic anisotropy which may help improve the diagnosis of various neurological diseases and disorders.

## 1 Introduction

Measuring and mapping nervous tissue microstructure non-invasively has been a long sought goal in neuroscience. Clinically, several neuropathologies such as cancer and stroke, are associated with changes in tissue microstructure. Over the past decades, significant MRI technical advances, including parallel imaging reconstruction, hardware and pulse sequence improvements, etc. all helped achieve millimeter voxel resolution. However, even at this length scale, neural tissue consists of highly heterogeneous and anisotropic micro-domains [1] despite its uniform appearance in conventional scalar valued MRIs. Diffusion tensor imaging (DTI) was able to resolve micron scale features, much smaller than the voxel size, [2, 3] by probing the diffusion of water molecules within a voxel modeled using a second order diffusion tensor estimated from single pulsed field gradient (sPFG) experiments. It was shown that the scalar diffusion weighting factor (i.e., b-value) in the Stejskal-Tanner experiment [4] needs to be replaced by a matrix or tensor (i.e., b-tensor). However, DTI provides only a mean diffusion tensor that is averaged over the entire MRI voxel, which belies the underlying heterogeneity of neural tissue [5, 6, 7]. Although some of these limitations could be overcome by increased spatial resolution, this comes at the cost of reduced signal-to-noise ratio (SNR) and increased scan time. The fundamental SNR limitation of MRI makes it impractical to resolve individual neuronal soma and axons whose sizes range between 0.1 - 60 *µ*m.

Multiple pulsed field gradient (mPFG) experiments have been shown to remove some degeneracies associated with DTI by differentiating between macro- and micro-scale anisotropy as demonstrated in yeast suspensions [8, 9], polydomain lyotropic liquid crystal systems [10], fixed gray matter [11] and in the live human brain [12, 13]. This has stimulated renewed interest in the diffusion MRI community to elucidate neural tissue microstructure beyond that provided by DTI [14, 15, 16]. However, many of these approaches generally rely on assumptions about the microstructure or morphology of the material, for instance that it consists of spherical and cylindrical pores, or of an ensemble of randomly oriented ellipsoids, limiting their applicability to complex neural tissue, especially pathological tissue where the changes in microstructure are not known.

A more general feature of neural tissue microstructure as observed in electron micrographs is a continuum of water compartments of various sizes, shapes and orientations, each surrounded by a plasma membrane. Although native MR imaging of these individual compartments is SNR prohibitive, imaging the statistical properties of an ensemble of these compartments within each voxel is feasible and may provide a sensitive measure of microstructural changes that occur during disease at the subvoxel scale. Given the permeability of water across the plasma membrane through the lipid bilayer, aquaporins, and transporter proteins [17], water diffusion within each microvoxel (i.e., the smallest compartment encoded in a diffusion experiment) may be treated as a hindered Gaussian diffusion process described by a single second-order diffusion tensor. Assuming the macroscopic MRI voxel is composed of a continuum of such non-exchanging microdomains or microvoxels at the time scale of the diffusion experiment, Basser et al. [18] proposed a probability density function in the space of diffusion tensors, also known as a diffusion tensor distribution (DTD), to describe such a microstructure. The resulting MR signal model was provided by Jian et al. as a linear superposition of Gaussian diffusion signal attenuation profiles arising from microvoxels [19]. Despite the use of single-PFG data by Jian et. al, it was later realized that mPFG data are required to unambiguously estimate the full DTD [20].

Estimating the DTD was shown to be an ill-posed inverse problem since it requires taking an inverse Laplace transform of the measured DWI [19]. Several approaches have been proposed in the literature to overcome the ill-posedness including using parametric distributions [19, 21], using the cumulant expansion of the MR signal to estimate the moments of DTD [20], and assuming symmetries of DTD [22, 23] with attendant limitations. In this study, we use a normal tensor-variate distribution (NTVD) whose samples are constrained to be positive definite (CNTVD) for DTD which we showed previously to be have useful properties such as maximum entropy, compactness and requiring only rank-1 and rank-2 b-tensors for their estimation, which overcomes some of the limitations of the previous approaches [24].

Given a theoretical framework for DTD estimation, implementing mPFG MRI experiments is challenging due to long echo times (TE) resulting from successive application of multiple gradient pairs, coherence transfer pathway artifacts from multiple refocusing radio frequency (RF) pulses [9]. Several single spin-echo variants of mPFG were introduced using combination of pulsed field gradients [25, 26, 27, 28] however these approaches may be cumbersome, as changing the shape of the b-tensor requires using a different set of gradients, and may suffer from concomitant gradient artifact known to bias DTD measurements [29]. A recent single spin-echo variant of mPFG involves using free optimized gradient waveforms [20, 30, 31] to generate arbitrary rank b-tensors and termed as *q*-trajectory imaging (QTI). Despite its generality, QTI has limitations such as a lack of welldefined diffusion time and diffusion pulse duration which limits the physical interpretation of the measured signals. The complex shaped gradient waveforms can be difficult to debug/implement. The asymmetry of the diffusion gradients around the 180^*◦*^ RF pulse despite being time efficient introduces concomitant gradient artifacts. This was partly overcome by symmetrizing the diffusion gradient around the 180^*◦*^ pulse [20], which increases TE, or by adding a constraint to their gradient shape optimization routine [29], which increases its complexity.

Here, we introduce a new versatile and easy to implement mPFG MRI experiment in a single spin echo without concomitant gradient artifacts, capable of generating arbitrary b-tensors of rank1, 2 or 3, using interfused gradient pulses. Our pulse sequence has well-defined diffusion gradient pulse widths, diffusion and mixing times, making it amenable to analysis and interpretation using the *q*-space formalism [32] with applications beyond DTD MRI. We used an optimized compressed sensing (CS) based experimental design that we proposed earlier [24] to uniformly sample b-tensors of varying shapes, sizes and orientations, which may be less biased compared to existing schemes. We disentangle the size, shape and orientation heterogeneity within a voxel and introduce a new method to perform tractography which naturally accounts for complex fiber configurations. We show that the derived heterogeneity measures and DTD tractography are capable of capturing established microstructural features and uncovering new ones in the human brain tissue *in vivo*.

## 2 Materials and Methods

### 2.1 Signal model

The MR signal from an ensemble of diffusion tensors distributed according to *p* (*D*_*ij*_) is given by [19],

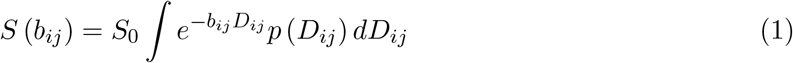

where *S*_0_ is the signal without diffusion weighting, and *b*_*ij*_, *D*_*ij*_ are the second order symmetric b-tensor and diffusion tensor respectively. Assuming a CNTVD for *p* (*D*_*ij*_), the signal equation is approximated using Monte Carlo (MC) integration with samples, *D*_*ij*_, drawn from a NTVD [18] with a given second-order mean, 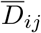, and fourth-order covariance tensor, Ω_*ijkl*_, which are filtered as shown below to ensure positive definiteness of the ensemble [24],

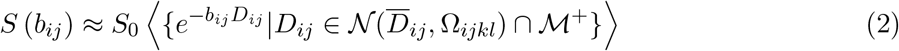

where the angle brackets denote ensemble averaging over all micro-diffusion tensors, 𝒩 is the NTVD and 𝔐^+^ is the space of positive definite second-order tensors. For convenience, the mean and covariance tensors of 𝒩 are expressed as a 6D vector and 6 *×* 6 symmetric positive definite matrix, respectively when drawing samples from the distribution.

### 2.2 MRI pulse sequence and experimental design

The new dPFG pulse sequence used to generate b-tensors of ranks 1 and 2 is shown in Figure 1 in comparison with the traditional and QTI methods. The two independent diffusion gradient pulses are incorporated within a single spin-echo sequence to reduce TE as compared to traditional dPFG sequence [9]. The use of PFG as opposed to free waveforms makes it easier to implement compared to QTI sequences. This pulse sequence design uses the waveform symmetrization strategy [33] to eliminate concomitant gradient artifacts, which worsen with increasing gradient strength and sample dimensions [34] and are particularly detrimental in DWI applications [29, 35]. The removal of concomitant gradient artifacts is accomplished by splitting the two gradient pairs of the preparation symmetrically on either side of the 180^*◦*^ RF pulse as shown in the figure.

**Figure 1:**
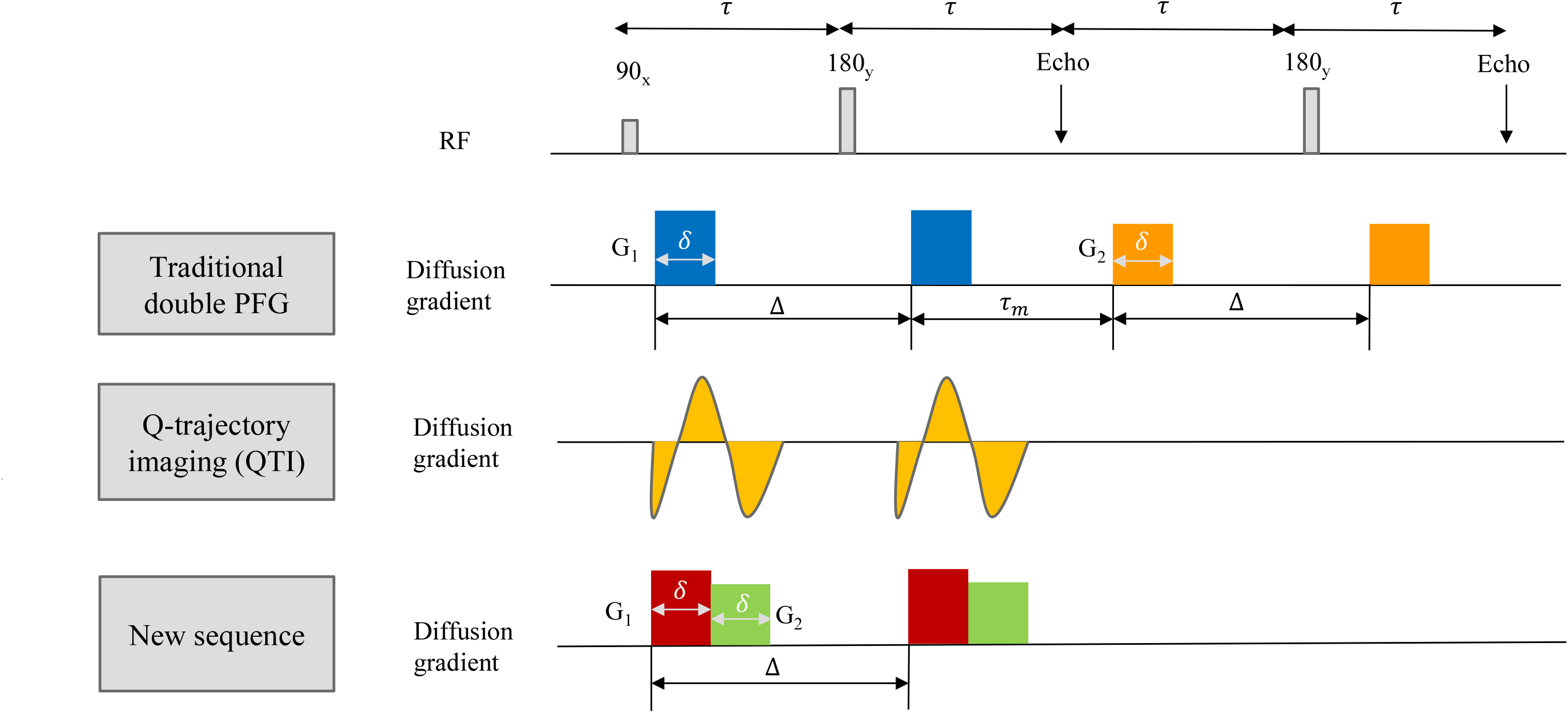
New interfused-pulsed field gradient (iPFG) for performing dPFG experiment in a single spin echo used to estimate the DTD compared with the traditional method and recent q-trajectory imaging (QTI). The new sequence combines the advantage of traditional and QTI method by performing the dPFG experiment in a single spin echo using standard pulsed field gradients capable of generating arbitrarily shaped rank-1 and rank-2 b-tensor while also immune to concomitant gradient and coherence artifacts. The two diffusion encoding gradient pairs, *G*_1_ and *G*_2_, may be applied along the same or different directions. The duration of the individual diffusion gradient pulses, *δ* and the diffusion time, Δ are also indicated.

A experimental design described in [24] used compressed sensing to generate a set of rank-1 and rank-2 b-tensors. Briefly, eigenvectors of the b-tensor were randomly rotated to uniformly sample orientation while their two non-zero eigenvalues were constrained so that their sums and ratios follow uniform distributions for size and shape, respectively. The diffusion gradient strengths/directions in the new pulse sequence required to generate the desired b-tensors for fixed *δ* and Δ are obtained using numerical optimization implemented in MATLAB (Mathworks, Natick, MA) constrained by the limits of the gradient hardware.

### 2.3 MRI measurements and pre-processing

MRI data was acquired using a 20-channel RF coil on a 3T scanner (Prisma, Siemens Healthineers) capable of up to 80 mT/m gradient strength and 200 T/m/s slew rate with echo planar imaging (EPI) readout. The measurement was performed on 24 and 47 year old healthy male volunteers who provided informed consent in accordance with a research protocol approved by the Institutional Review Board (IRB) of the Intramural Research Program of the National Institute of Neurological Disorders and Stroke (NINDS). To test reproducibility of the DTD estimation pipeline, one of the volunteers was scanned twice in separate imaging sessions. The relevant acquisition parameters are as follows: *δ\*Δ = 14*\*40 ms, TR*\*TE = 3500*\*90 ms, field of view (FOV) = 210 mm *×* 210 mm *×* 150 mm, 1.5 mm in-plane spatial resolution at 1490 Hz/pixel bandwidth, and 5 mm thick slices with GRAPPA acceleration factor = 3 and partial Fourier = 6/8. A total of 216 different b-tensors of uniform size, shape and orientation shown in Figure 2 were sampled with b-values ranging from 0 - 2000 s/mm^2^. The b = 0 s/mm^2^ acquisition was repeated with reversed phase encoding direction to correct for susceptibility induced geometric distortions.

**Figure 2:**
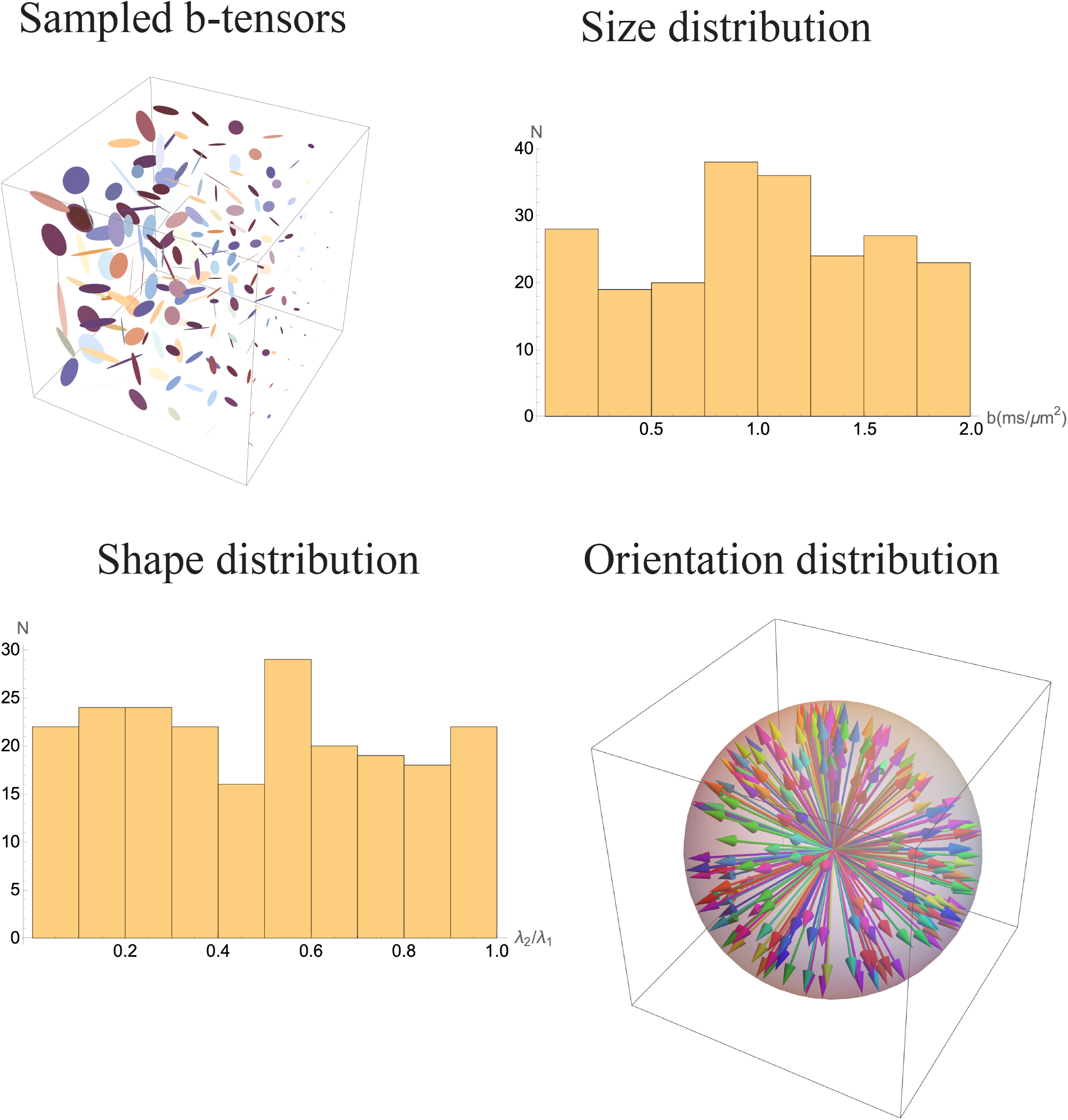
Experimental design for DTD MRI using 216 diffusion encoding rank-1 and rank-2 btensors depicted by ellipsoids (top left). The uniform size and shape distribution of b-tensors are shown using the histogram of their b-values (top right), and the ratio of the two non-zero eigenvalues (bottom left) respectively. The uniform orientation distribution of b-tensors are shown by plotting the ensemble of their primary eigenvectors (bottom right).

The individual DWIs were processed in the following order prior to DTD reconstruction, 1) thermal noise was suppressed using a Marchenko-Pastur principal component analysis (PCA) algorithm [36] implemented in the DIPY software environment [37], 2) the effect of inter-scan subject motion and eddy current induced geometric distortion were reduced by registering the DWIs with b = 0 s/mm^2^ acquisition using 3D affine transform [38, 39] implemented in FSL software [40], and 3) geometric distortions arising from inhomogeneities in the static field were accounted for using the off-resonance magnetic field estimated from b = 0 s/mm^2^ acquired with reverse phase encoding implemented in FSL software environment [40, 41].

### 2.4 Estimation of microstructural parameters and tractography

The mean and covariance tensors characterizing the CNTVD were both estimated from the MR signals using methods outlined in [24] except that Cholesky decomposition is now used to impose the positive definiteness constraint of the covariance tensor which improves the signal fit. Briefly, the different symmetries of the mean [42] and covariance tensors [43] were exploited to build a family of nested models from which the most parsimonious one was chosen using the Bayesian information criterion (BIC). For a given nested model, the MR signal is fit to Equation (2) using simplex-type numerical optimization algorithm to estimate the unknown parameters.

The estimated CNTVD parameters are used to delineate several microstructural features within the voxel, which are also described in greater detail in [24]. Briefly, the micro-diffusion tensors in the voxel are simulated by drawing MC samples from the CNTVD with the estimated mean and covariance tensors. The macroscopic fractional anisotropy (FA) and macroscopic orientation distribution function (ODF) are computed from the mean diffusion tensor, whereas the microscopic FA and microscopic ODF (i.e. *µ*FA and *µ*ODF, respectively) are computed by an ensemble average of the FA and ODF, respectively, obtained by summing over each of the micro-diffusion tensors. The “macro” and “micro” quantities are generally not equal due to the non-commutativity of the associated operators (e.g. FA, ODF, etc).

The tissue heterogeneity is classified based on the size, shape and orientation heterogeneity of the micro-diffusion tensors. The size and shape of micro-diffusion tensors are quantified by the trace (i.e. mean diffusivity, MD) and FA-weighted eigenvalue skewness of the diffusion tensor [44] respectively, whose distribution are measured and mapped. The orientation heterogeneity is quantified by *V*_orient_ which is the extent of the orientation dispersion of micro-diffusion tensors [24]. Further, tractography was performed on the whole brain using the macro and micro ODFs. For better delineation of ODF peaks, ODFs were filtered/sharpened using the deconvolution transform implemented in DIPY software environment [37] prior to performing tractography [45]. Streamline tractography was then performed using the filtered macro and micro ODFs in MRTrix software environment [46].

## 3 Results

Heterogeneity and microscopic anisotropy were observed in varying degrees in all regions of the brain except the corpus callosum where the anisotropic DTI model was chosen by the model selection pipeline with zero covariance. Results from a representative slice in cerebellum and cerebrum are provided in this section to show their agreement with known findings and the new information provided by our DTD framework over existing diffusion signal models for brain tissue. Reproducibility of DTD measures was addressed in a “test-retest” paradigm by comparing them at approximately the same slice in a twice scanned subject.

The size heterogeneity quantified by the spectral moments of the mean diffusivity are shown in Figure 3. The mean was uniform in the parenchyma, however the higher moments such as standard deviation and skewness showed new contrast. In particular, the skewness was elevated in cerebellar and sub-cortical gray matter highlighted using red and pink arrows in the figure, respectively. In CSF-filled regions, the standard deviation was uniform and elevated while the skewness was lower inside the ventricles (≈ 0.4) compared to that at the ventricle-tissue boundary (≈ 0.9).

**Figure 3:**
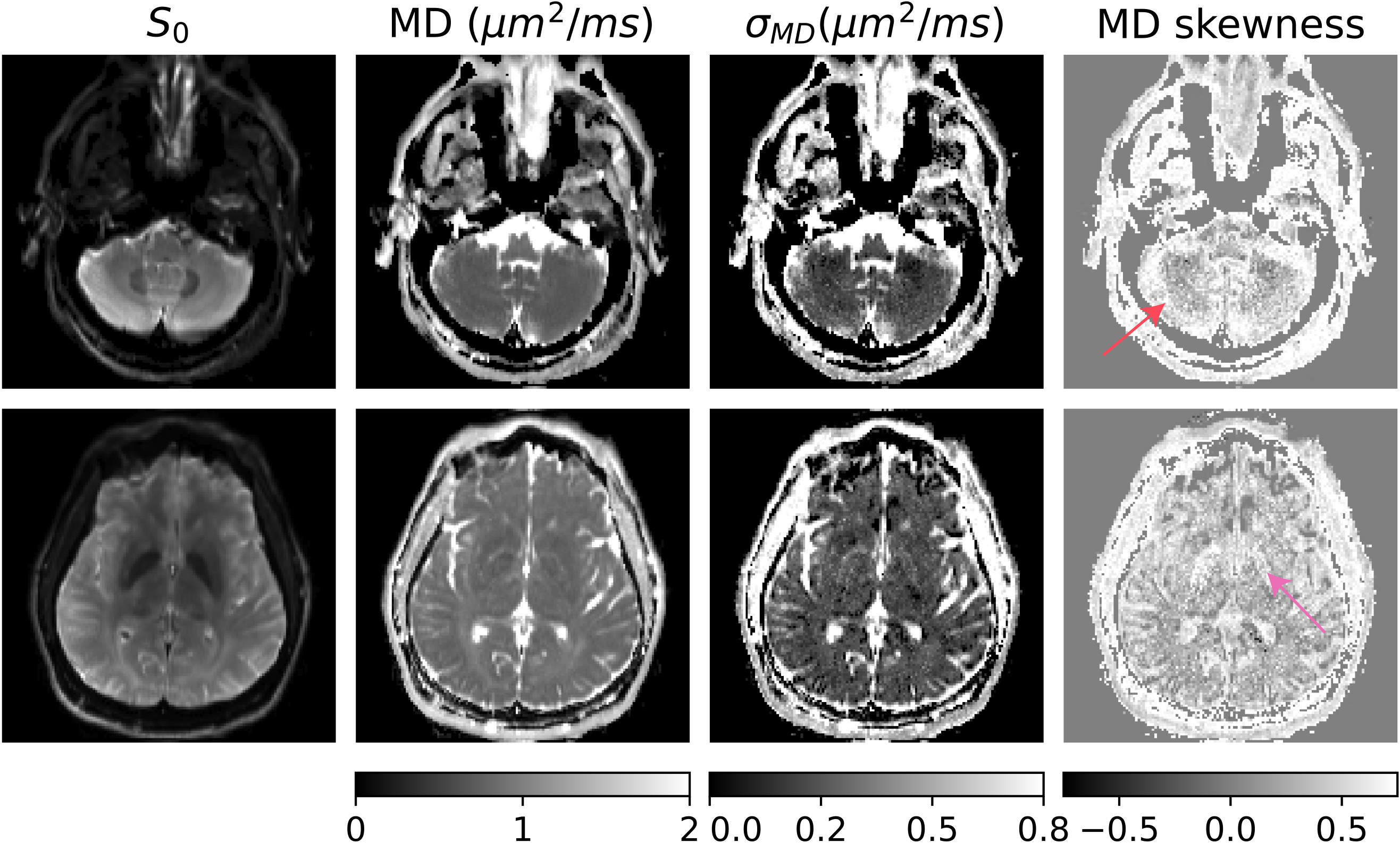
Size heterogeneity in the brain and cerebellum quantified by the moments of the mean diffusivity (MD) spectrum in two different slices from a representative subject. The maps of mean, standard deviation (*σ*), and skewness of the MD spectrum are provided along with *S*_0_ map for reference. Spectral moments beyond the mean revealed new contrast in cerebellar and cerebral gray matter shown using arrows in the skewness map despite the mean being uniform in the parenchyma.

The shape heterogeneity in the brain given by the moments of the shape measure is shown in Figure 4. The shape of the mean diffusion tensor within the slice was overall positive (i.e. ranging from spherical to prolate) with higher values in coherent white matter regions compared to that in gray matter and complex fiber regions. The standard deviation was uniform with high values (≈ 0.4) in parenchyma compared to that in CSF filled regions (≈ 0.2). The skewness was negative overall in coherent white matter while positive in other areas. The cerebellar gray matter showed layered structure in the skewness map shown using red arrow in the figure which was not evident in the mean. The sub-cortical gray matter highlighted in the size heterogeneity measure also showed elevated shape heterogeneity measure shown using pink arrow in the figure.

**Figure 4:**
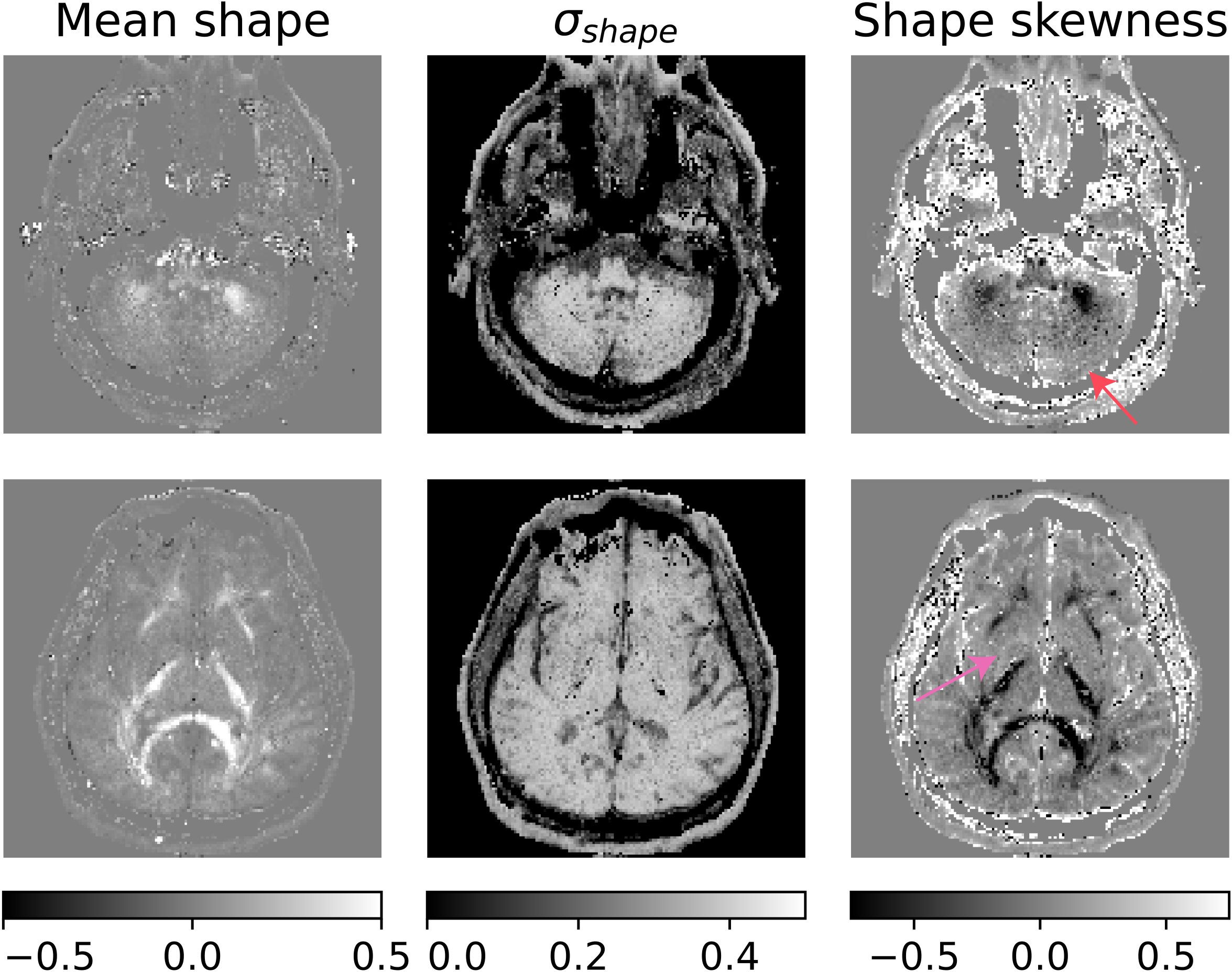
Shape heterogeneity in the brain and cerebellum quantified by the spectral moments of the shape measure defined by the FA-weighted eigenvalue skewness of the diffusion tensor. Spectral moments beyond mean showed new contrast such as the layers in cerebellum (red arrow), and within sub-cortical gray matter (pink arrow).

The orientation heterogeneity in the brain explained using *µ*FA and *V*_*orient*_ measures are shown in Figure 5. The FA and color-coded primary eigenvector of the mean diffusion tensor (direction-encoded color, DEC) are also provided for reference. The DTI based FA and DEC map showed orientation configuration of white matter tracts consistent with previous findings. The *µ*FA map was equal to FA in highly coherent white matter tracts such as the corpus callosum while elevated in gray matter and complex fiber regions in white matter such as in the corona radiata shown using a green arrow in the figure. It was smallest in CSF-filled ventricles and CSF-tissue boundaries. The *V*_*orient*_ measure was approximately inversely proportional to FA with large values observed in the cerebral cortex and cerebellar gray matter which contributed to the increased *µ*FA observed in these regions.

**Figure 5:**
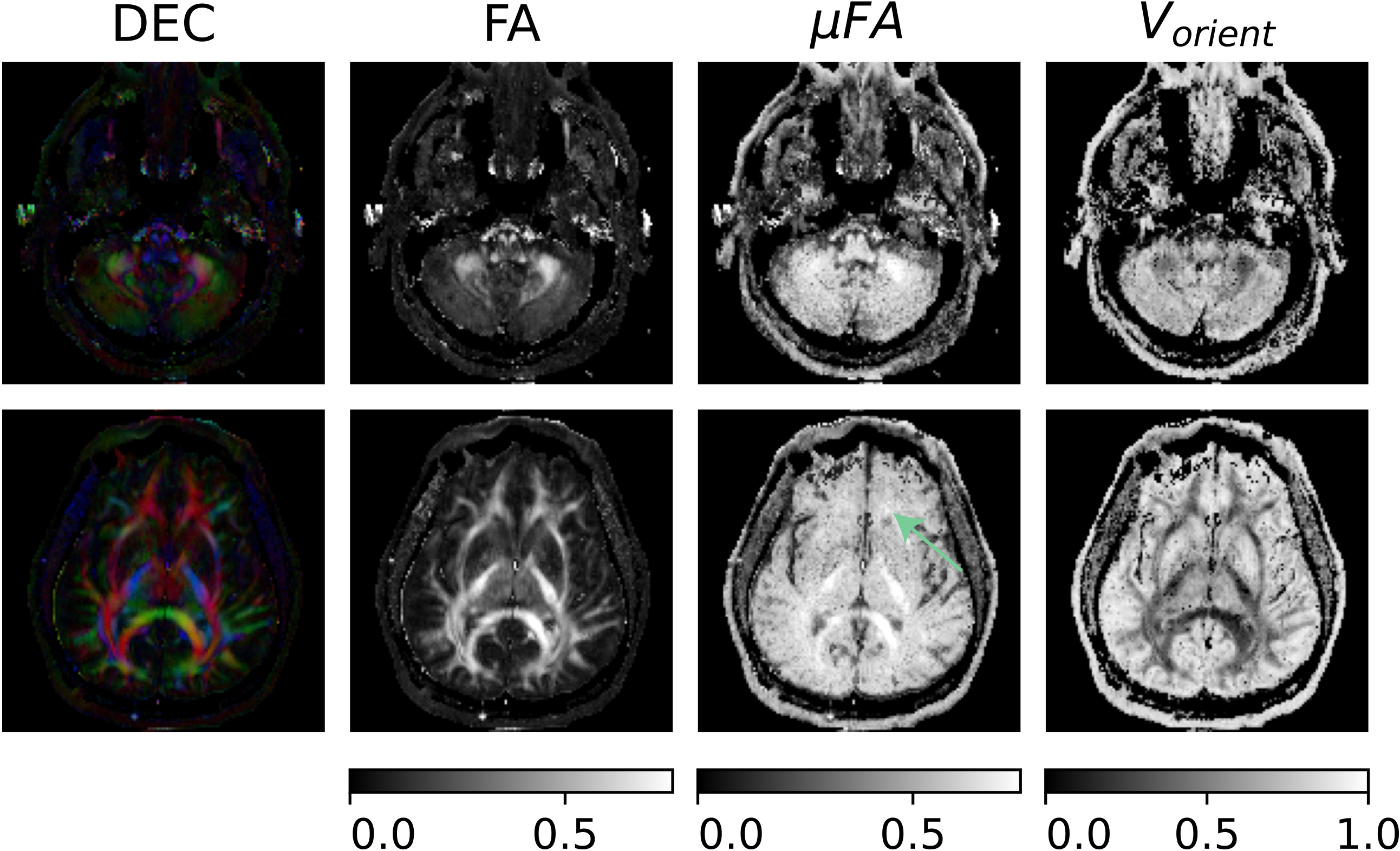
Orientation heterogeneity in the brain and cerbellum quantified by the fractional anisotropy (FA), microscopic FA (*µ*FA), and *V*_*orient*_ measures. The color coded primary eigenvector of the mean diffusion tensor (direction encoded color, DEC) is provided for reference. The *µ*FA was uniform and elevated in gray matter compared FA in the region shown for example using the green arrow. *V*_*orient*_ was higher in coherent white matter fiber tracts and vice-versa which explains the *µ*FA observed in these regions.

In addition to providing new contrast, DTD MRI helps resolve confounds associated with DTI. For example, the reduction in FA observed in corona radiata (green arrow in Figure 5) could be a result of edema that may occur in disease or due to microscopic anisotropy arising from orientation or shape heterogeneity. DTD MRI measures could help disentangle this ambiguity by indicating that the dominant heterogeneity in this region is due to that of shape as *σ*_*MD*_ and *V*_orient_ are both small. Thus the FA loss in the region of interest (ROI) is not due to edema, as it would have resulted in increased *σ*_*MD*_, but rather resulting from high shape heterogeneity in the region which might be due to splaying of fibers at sub-microvoxel scale.

The reproducibility of DTD measures is demonstrated by comparing the results obtained on the same subject scanned twice in separate imaging sessions at approximately the same slice location as shown in Figure 6. The results are also compared quantitatively by averaging the DTD measures over ROIs encompassing portions of gray matter, white matter and CSF which are tabulated in Table 1. It can be observed that several spatial features of the measures were recaptured on both the repetitions. This includes the increased *σ*_*MD*_ observed at the CSF filled regions, reduced MD skewness inside the ventricles, uniform *σ*_*shape*_ in the parenchyma and negative shape skewness in coherent white matter, and reduced *V*_*orient*_ in coherent white matter and vice-versa. This is also quantitatively reflected in similar averaged values of these measures in the gray matter, white matter and CSF ROIs as shown in the table.

**Table 1:**
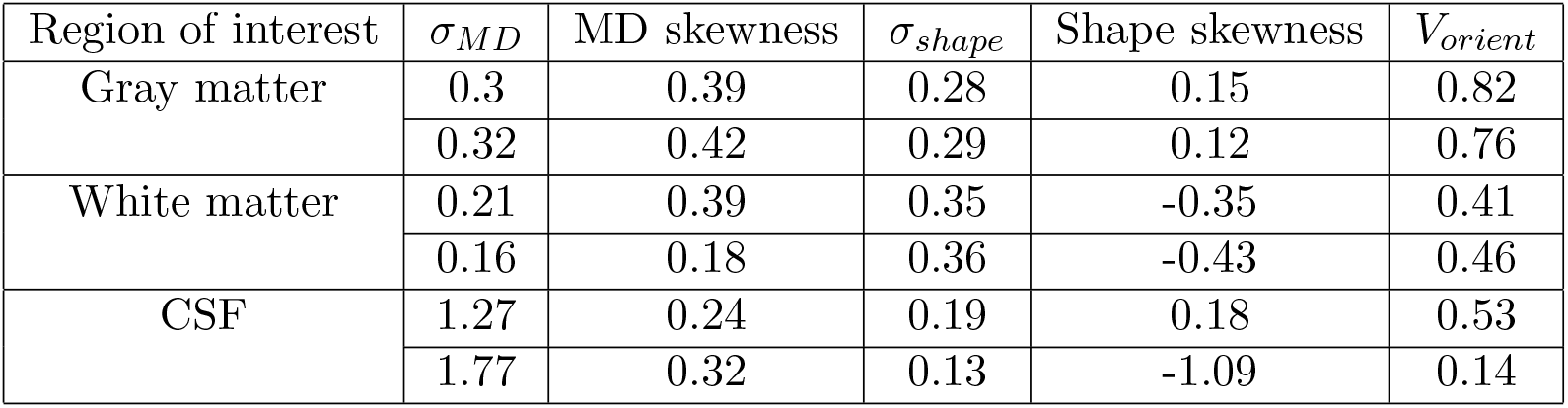
Reproducibility of the DTD analysis demonstrated by averaging the value of measures over gray matter, white matter and CSF regions of interest (ROI) on data acquired on the same subject on two different sessions (shown in rows) within each ROI. Gray matter ROI was chosen over the cortex, white matter ROI covers a region of the internal capsule, and CSF ROI was drawn over a part of the lateral ventricles. It can be observed that the values of these measures match despite slight changes in the excited slice.

| Region of interest | $\sigma_{MD}$ | MD skewness | $\sigma_{shape}$ | Shape skewness | $V_{orient}$ |
| --- | --- | --- | --- | --- | --- |
| Gray matter | 0.3 | 0.39 | 0.28 | 0.15 | 0.82 |
|  | 0.32 | 0.42 | 0.29 | 0.12 | 0.76 |
| White matter | 0.21 | 0.39 | 0.35 | -0.35 | 0.41 |
|  | 0.16 | 0.18 | 0.36 | -0.43 | 0.46 |
| CSF | 1.27 | 0.24 | 0.19 | 0.18 | 0.53 |
|  | 1.77 | 0.32 | 0.13 | -1.09 | 0.14 |

**Figure 6:**
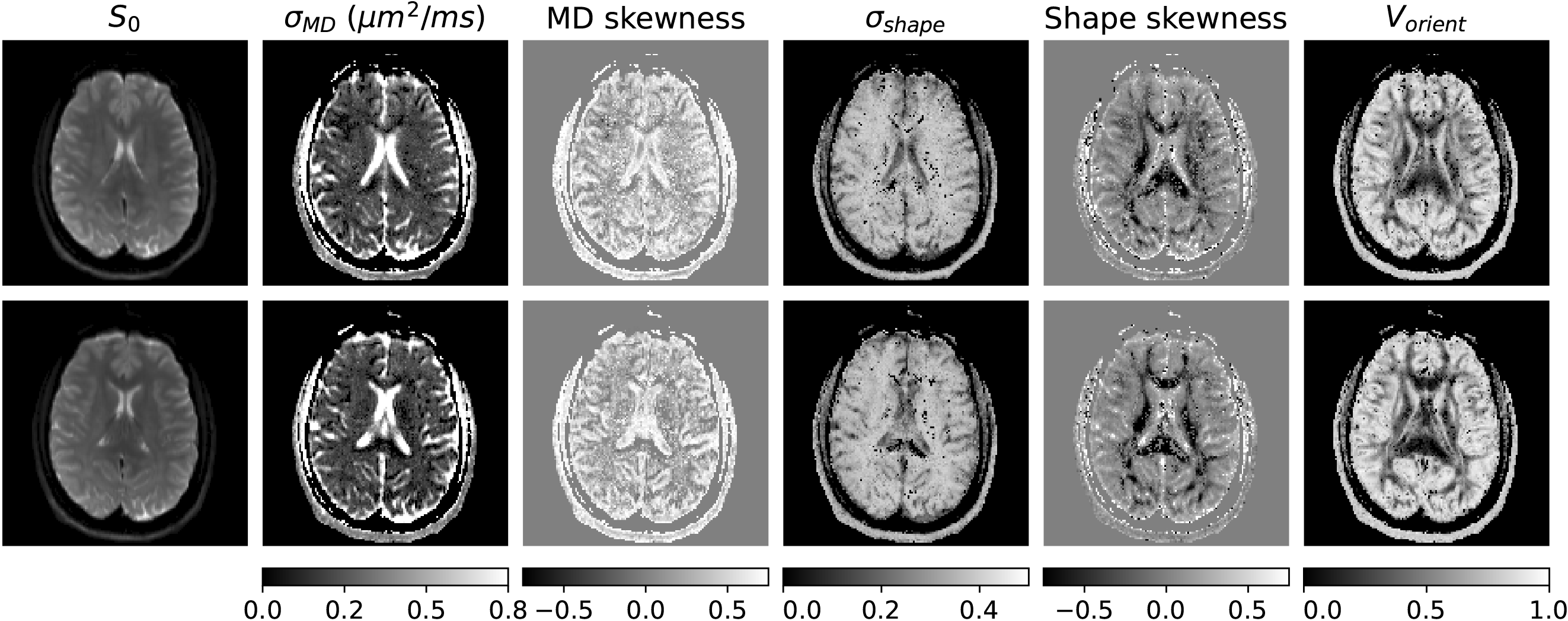
Test-retest of DTD analysis demonstrated by scanning the same subject twice in separate sessions. DTD derived measures at approximately the same slice location in both the runs (top and bottom rows) are shown along with *S*_0_-map provided for reference. The high *σ*_*MD*_ in ventricles, uniform *σ*_*shape*_ in the parenchyma, and correspondence between *V*_*orient*_ and coherence of fiber tract regions are captured on both scans.

The orientation dispersion in brain tissue is shown by comparing the macro and micro ODFs in a few ROIs shown in Figure 7. The ROIs were chosen in corpus callosum, corona radiata and internal capsule fibers. In the corpus callosum ROI, the macro and micro ODFs were identical. In the corona radiata ROI, macro ODFs showed a single fiber population while the *µ*ODFs clearly showed kissing and/or crossing of two fiber populations. In the internal capsule ROI, the macro ODFs reflected two fiber populations with unique principal direction as shown by different colors in the DEC map. The *µ*ODFs however clearly showed almost 90-degree crossing between the two fiber populations especially at the interface.

**Figure 7:**
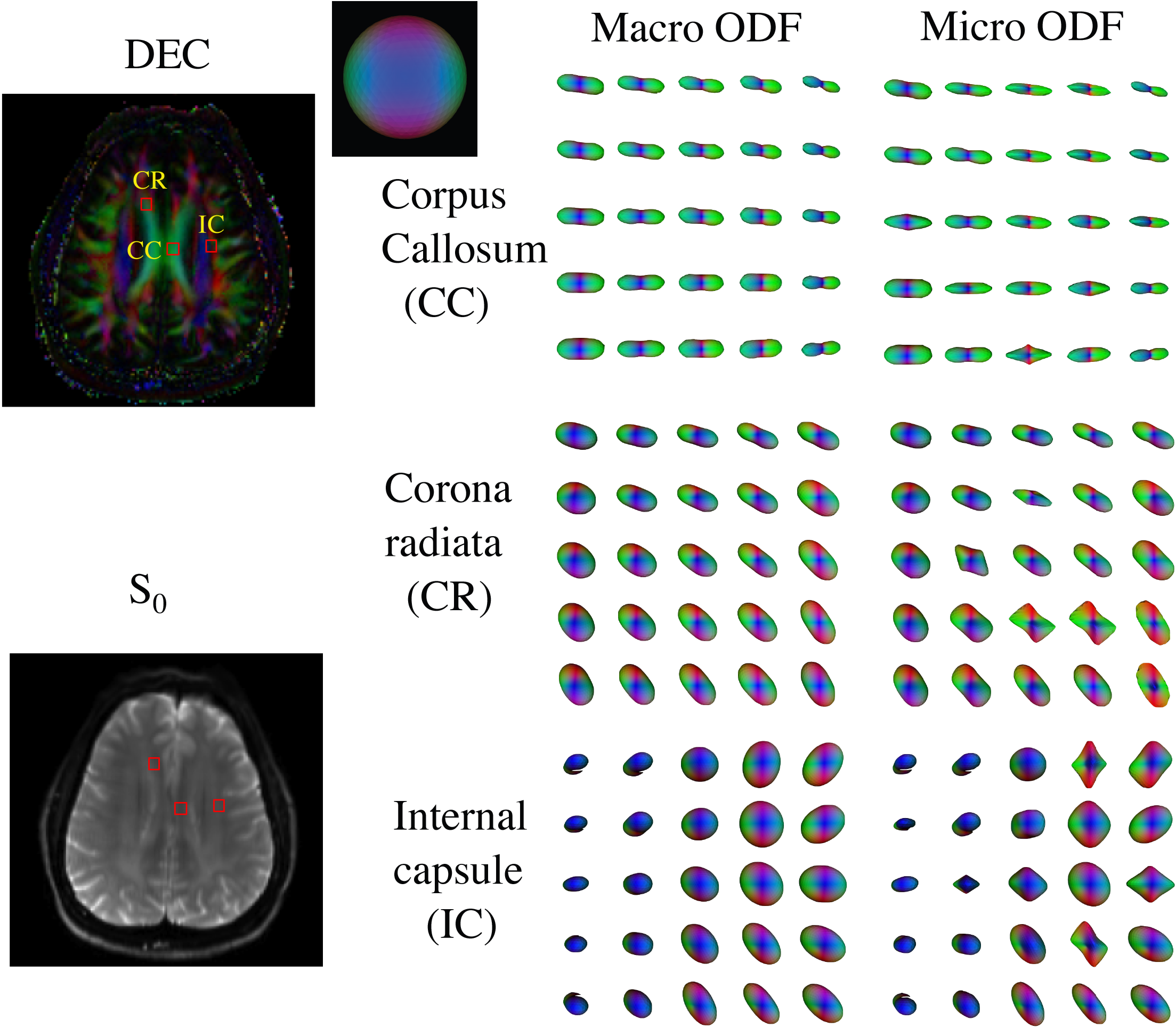
Orientation dispersion in brain tissue described using macro- and micro-orientation distribution functions (ODF) in the following three brain ROIs: 1) corpus callosum (CC) showing identical macro and micro-ODFs, 2) corona radiata (CR) with *µ*ODFs showing splaying fibers, and 3) internal capsule (IC) with *µ*ODFs showing crossing fibers but the macro-ODF showing more spherical profiles. The chosen ROIs are displayed with direction-encoded color (DEC) map obtained from the primary eigenvector of the mean diffusion tensor and *S*_0_-map for reference.

The results of streamline tractography performed for the whole brain using macro and micro-ODFs are shown in Figure 8 for a coronal slice. DTI tractography captures the fibers accurately in coherent white matter regions such as the body of corpus callosum, as expected. Differences between DTI and DTD tracts were observed in several white matter regions with complex fiber configurations and are highlighted in the figure using arrows. The projections of corpus callosum fibers to multiple areas of cerebral cortex absent in DTI tractography were clearly visible in DTD tractography.

**Figure 8:**
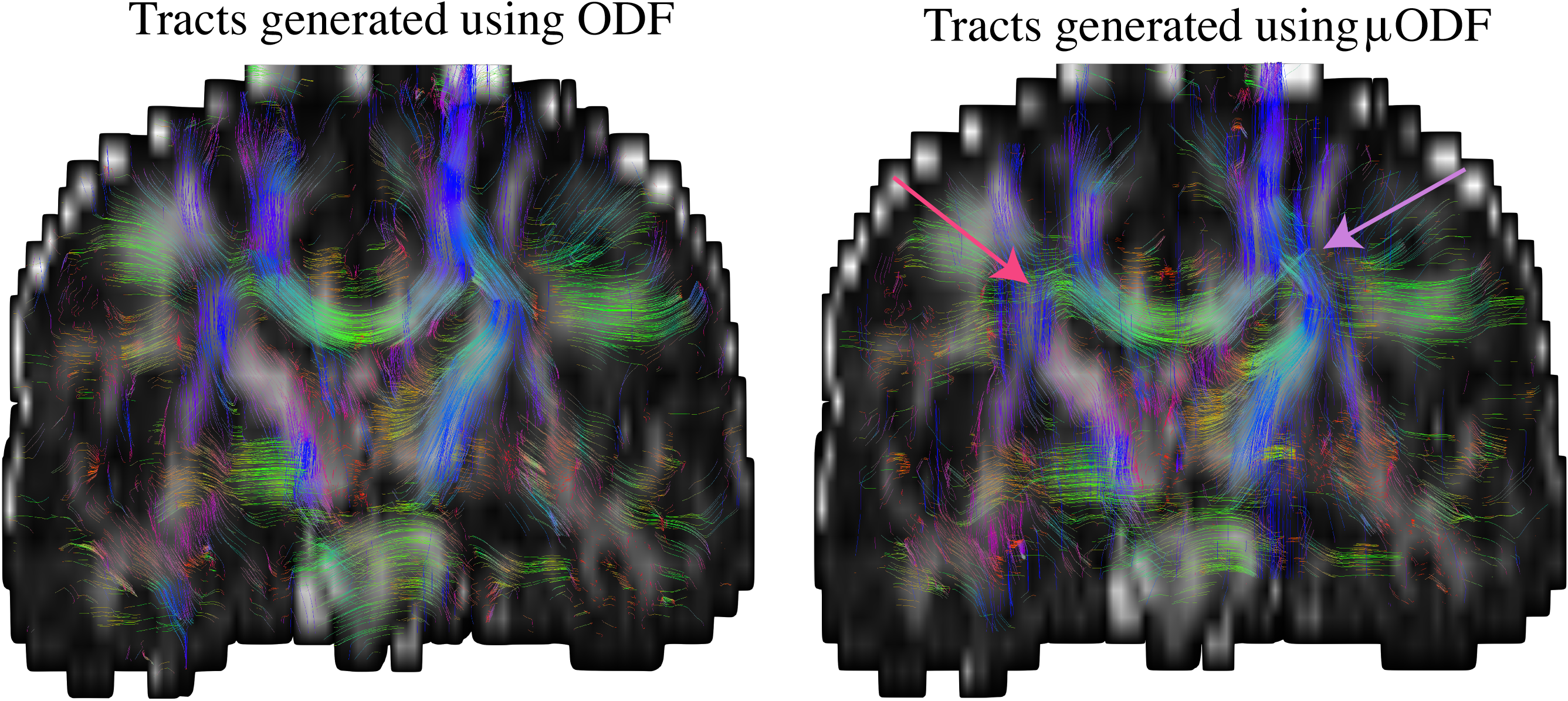
Streamline tractography using macro or DTI-ODFs and micro-ODFs on a coronal slice with the FA map overlaid in the background for reference. The macro ODF is derived from the mean diffusion tensor while the micro-ODF is derived from the individual tensors constituting the DTD (Left) Tracts obtained using DTI ODF. (Right) Tracts obtained using DTD *µ*ODF showing multiple crossing fibers which are absent in the tracts generated from the DTI ODF. A few crossing fiber regions resolved by *µ*ODF based tracts are shown using arrows.

## 4 Discussion

### 4.1 A new paradigm for mPFG experiments

The source of non-Gaussian like diffusion of water observed in neural tissue could be either due to restriction or heterogeneity as both produce a similar effect on the MR signal. This had led to a proliferation of models for neural tissue microstructure which either assume restriction [47, 48] or heterogeneity [19, 25, 49] or both [50, 51]. The applicability of these models strongly depend on relationship between the timing parameters of the diffusion gradient pulses and the pore/compartment sizes. While this is often the case for traditional mPFG or multiple diffusion encoded (MDE) experiments, it is not the case for the recent clinically viable adaptations, such as QTI, which uses complex gradient waveforms. Our iPFG MRI sequence on the other hand has well defined diffusion gradient parameters which enables us to analyze our signal decay data using either of the two modeling frameworks. The mPFG nature of our experiment is shown in Section 1 in the supporting material where the b-tensor integral is analytically calculated for our gradient pattern ignoring the ramps and shown to possess a rank of 2.

By extending the existing *q*-space formalism to iPFG, we show that our sequence retains several salient features of traditional mPFG experiments [32] thus showing its versatility for use in applications beyond DTD MRI. The echo attenuation, *E*, resulting from an ensemble of compartments is given by,

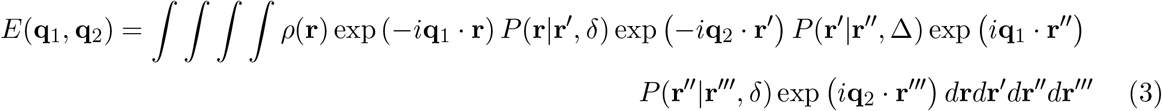

where **q** is the time integral under the diffusion gradient pulse, **r** is the spin position, *ρ*(**r**) is the initial spin density at **r** and *P* (**r**_1_|**r**_2_, *dt*) is the propagator which is the conditional probability for the spins to move from **r**_1_ to **r**_2_ in time interval, *dt*.

Considering restricted diffusion within the pores, the above expression is evaluated for two time regimes. In the limit the spins have traversed the entire pore during the mixing time (*τ*_*m*_ = *δ*), the propagator becomes equal to the spin density. The resulting expression for echo attenuation is identical to that obtained from the traditional dPFG analysis as shown below (derived in Section 2.1 in the supplementary material),

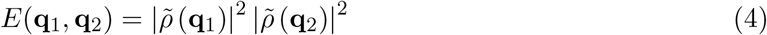

where 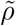 is the Fourier transform of the spin density, also known as the pore spectral density. In the limit of short diffusion time, the propagator reduces to a delta function resulting in the following expression for the echo attenuation derived in Section 2.2 in the supplementary material,

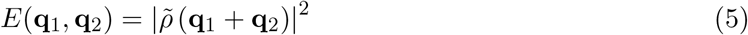

which is different from that obtained from a traditional dPFG analysis [32]. However, expanding the pore spectral density using a Taylor series, it can be shown that the new method retains salient features of earlier dPFG experiments such as the angular dependence of the signal with relative orientation of the two gradient pairs [32] derived in Section 2.2 in the supplementary material. It also shows that the diffusion gradient parameters that determine the sensitivity of the measurement to microstructural features are well defined in our measurement unlike the complex *q*-trajectory or waveform approaches.

### 4.2 A new form of tractography using DTD MRI

Since the introduction of DTI tractography [52, 53], a slew of methods were introduced to overcome its limitations resulting from complex fiber configurations in white matter [54, 55, 56, 57]. A common feature of these methods is that they all use sPFG data to reconstruct fiber tracts. While sPFG acquisitions are sufficient for reconstructing fiber tracts with highly homogeneous subvoxel microstructure, mPFG measurements encode correlations between diffusion tensors and may therefore be better suited to disentangle fiber populations with different diffusion properties within the voxel. However, translating existing fiber tracking methodologies to mPFG data is partly hampered by diffusion encoding using multiple *q*-vectors. Recently this was partly overcome by replacing multiple *q*-vectors with the symmetry axis of the b-tensors while performing spherical deconvolution to estimate the fiber ODF [58] which is phenomenological. In addition, the limitations and challenges of performing spherical deconvolution are well known [59].

In this study, we introduced a new way to perform tractography by estimating the DTD in each voxel and deriving the corresponding *µ*ODFs from it. This is the first time a continuous mixture model is used to perform tractography on mPFG data with the added advantage of adaptability to any form of DTD not restricted to CNTVD. The DTD-derived tracts clearly demonstrate the ability to resolve multiple distinct fiber populations compared to DTI derived macro ODF despite the large slice thickness employed in this study needed for adequate SNR.

### 4.3 Sources of variability in DTD MRI

We assume the variability in the diffusion tensors within a voxel arises purely from the underlying tissue microstructure however, there are other sources of variability, as well. These include thermal noise, bulk motion of the sample, susceptibility induced gradients causing geometric distortion, and gradient hardware imperfections such as eddy currents, concomitant gradient fields, etc., which also result in the MR signal deviating from the DTI model. These effects may be particularly pronounced in clinical scanners.

We have partially accounted for bulk motion of the sample and eddy currents by registering the DWIs with a non-diffusion weighted volume, and susceptibility artifacts using a field map based correction. We have reduced the effect of random thermal noise using an existing Marchenko-Pastur PCA algorithm and our novel model selection pipeline given that the noise is independent and uncorrelated to tissue variability. We have further increased the SNR by shortening the TE using pulsed diffusion gradients, and eliminated concomitant gradient field dependent signal loss altogether. The agreement between our results to known brain anatomy and their reproducibility *in vivo* demonstrate the validity of our model in describing the underlying tissue microstructure, and robustness of our estimation pipeline.

### 4.4 Biophysical basis for DTD in neural tissue

The Gaussian micro-diffusion compartments observed in neural tissue could arise from the heterogeneous geometry of neuronal soma, axons, and extra-cellular spaces. In gray matter, sources of size and shape heterogeneity include known variability in the size and shape of neurons while the orientation heterogeneity could arise from a powder-averaged distribution of cells [60]. In white matter, size heterogeneity could result from differences in axon diameters. The shape heterogeneity could originate from complex fiber configurations known to exist on a sub micro-voxel scale [61], which would result in a mixture of diffusion tensor shapes within a voxel. The orientation heterogeneity could arise from kissing/crossing/splaying/merging of fiber tracts on the scale of the micro-voxel. In addition, presence of free water regions [62] with higher MD could introduce size and shape heterogeneity in both gray and white matter.

In the central nervous system (CNS), the neuronal soma diameters range from 10 - 50 *µm*, and axon diameters range from 0.16 - 9 *µm* with the mean diameter less than 1 *µm* [63]. The gaps in the extracellular space in the neural tissue range from 38 - 64 nm [64]. The ability of DTD MRI to capture tissue heterogeneity at these length scales depends on three conditions, 1) the net displacement of spins during the application of the gradient (i.e., 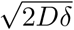) should be less than the size of the compartment to preserve its unique label [65], 2) the pitch of the magnetization phase helix wound on the sample by the diffusion gradient should be sufficient to cause an appreciable signal decay as the spins explore the compartment (i.e., pore size 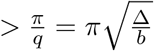 assuming half-pitch is sufficient), 3) the residence time of spins within the compartment should be longer than Δ which would otherwise result in averaging of compartments. Thus sensitizing small compartments requires rapid slew rates, large gradient strengths and short diffusion times. We believe these requirements make DTD-MRI particularly well-suited to implementation on high performance gradients such as [66, 67, 68].

Given the diffusion gradient parameters and MD of the brain, the resolution limits imposed by conditions 1 and 2 are approximately between 5 and 15*µm*, respectively. The diffusion time in our measurement averages compartments exchanging at a rate faster than 25*s*^*−*1^. While there is no clear relationship yet between exchange rate and length scale in neural tissue, it can be expected that the exchange rate scales with surface to volume ratio (SVR), and thus is inversely related to size, and directly related to water permeability of the compartment. Water exchange rate in healthy neural tissue measured using NMR is reported to be approximately 2*s*^*−*1^ [69, 70] which is slower than that imposed by our measurement. However values as high as 100*s*^*−*1^ have been recently found in live mouse spinal cord using diffusion exchange measurements with very strong static gradients that sensitize it to micron scale structures, thus posing a challenge for DTD MRI [71]. Together the effective micro-voxel size we are probing using this modeling framework and our experimental design for live human brain is approximately 15 *µm*. At this length scale, it is clear that individual axons in white matter and extra-cellular space cannot be resolved by our measurement, which is consistent with our finding that highly aligned fibers in the corpus callosum are adequately described by the DTI model (with a single mean tensor and a zero covariance tensor or no variability). However, heterogeneity in bundles of axons on a scale larger than 15 *µm* such as in kissing/crossing/splaying/merging fiber tracts should be detected by our measurement, which we do observe in regions such as the corona radiata and internal capsule.

### 4.5 Microstructural information from DTD-MRI

A key advantage of using CNTVD is to obtain the entire DTD since non-parametric estimation of the DTD is difficult. This statistical DTD model allows us to synthesize an arbitrarily large number of micro-diffusion tensors, which is not done in parametric or analytical models such as those based on cumulants or kurtosis, respectively. Such statistical information is required for accurately estimating salient microscopic quantities such as size and shape spectra of micro diffusion tensors and *µ*ODF that help reveal new structural features. By comparing our results with known anatomy, we show that the CNTVD may be sufficient to accurately capture key features of brain tissue microstructure. The higher *σ*_*MD*_ in ventricles and subarachnoid spaces is likely due to pulsatile flow of CSF from cardiac and respiratory motion previously observed *in vivo* [72].The small MD skewness observed in the ventricles indicates a Gaussian distribution of tensors expected in a free water region, while the higher skewness in the CSF-tissue boundaries might be due to mixing of the two compartments with very different diffusivities [25, 73]. The lower *σ*_*shape*_ measure in CSF-filled ventricles is consistent with the spherical distribution of tensors in this region. The negative shape skewness in coherent white matter may be due to the preponderance of prolate diffusion tensors in this region. The increased *µ*FA with respect to the macroFA in complex fiber regions is in agreement with that previously reported in the literature [74]. The dependence of *V*_*orient*_ measure with the degree of coherence of fiber bundles is as expected, and its non-zero value in most of the white matter regions is in agreement with 3D rendering of electron micrographs of white matter showing complex fiber configurations [61]. The kissing/crossing of fibers in the corona radiata observed in the *µ*ODF is well known as is the 90^*◦*^ crossing in internal capsule fibers [75].

The cerebellar gray matter region has been investigated in a previous study using spherical tensor encoding [76] where it was hypothesized that the signal persisting at high b-values results from the so-called “dot-compartment” with a single apparent diffusivity. The positive MD skewness in this region revealed by DTD shows that a persistent signal may instead arise from a multitude of small cells having varying diffusivities. The layered structure observed in the skewness map might have resulted from CSF contamination at the boundaries. In the cerebrum, the bilateral increase in MD and shape skewness in the subcortical gray matter region might be reflective of the diverse cell size and shapes in this region. It can be expected that new information could be further gleaned when studying pathological conditions. For example, inflammation in white matter would be reflected as increased *σ*_*MD*_ and concomitant reduction in shape skewness due to the presence of more spherical micro-compartments.

## 5 Conclusion

In this study, we present a new suite of tools to measure and map the DTD in live human brain. We introduce a new method to perform mPFG MRI experiments in a single spin echo sequence capable of generating b-tensors of ranks one, two, or three, without concomitant gradient artifacts. We demonstrate that our experimental design and signal inversion framework is able to capture heterogeneity in neural tissue. However, one of the limitations of our Monte Carlo approach is the long computation time that can be overcome in the future by performing the model fitting using graphical processing units (GPU).

It should be noted that even at the spatial resolution and with the diffusion gradients used in this study, macroscopic heterogeneity may dominate in certain voxels especially those with CSF such as in the cerebral cortex. With the advent of advanced image readout strategies and stronger gradients [66, 67, 68], it may be possible to achieve true mesoscopic resolution in a greater proportion of voxels by performing DTD at high *k*- and *q*-space resolution. This would help advance the application of our new heterogeneity measures and tractography in assessing new disease, normal and abnormal developmental processes, degeneration and trauma in the brain and in other soft tissues.

## 6 Acknowledgments

We thank Evren Özarslan and Michal E. Komlosh for thoughtful discussions and feedback. We thank Sinisa Pajevic for helping accelerate the DTD programming pipeline. We thank Wolfgang Resch for assisting with computations performed on the NIH HPC Biowulf cluster. This study was supported by the Intramural Research Program of the NICHD and NINDS. This work was also partly funded by NIH BRAIN Initiative grant 1U01EB026996-01 - “Connectome 2.0: Developing the next generation human MRI scanner for bridging studies of the micro-, meso- and macro-connectome.” and the CNRM Neuroradiology/Neuropathology Correlation/Integration Core, 309698-4.01-65310, (CNRM-89-9921). This work utilized computational resources of the NIH HPC Biowulf cluster (http://hpc.nih.gov). The opinions expressed herein are those of the authors and are not necessarily representative of those of the Uniformed Services University of the Health Sciences (USUHS), the Department of Defense (DOD), the NIH or any other US government agency, or the Henry M. Jackson Foundation for the Advancement of Military Medicine, Inc.

## Author Contributions

KNM, DG and PJB designed the research. KNM wrote the pulse sequence, performed the measurement and DTD analysis, and drafted the manuscript. AVA and JS assisted with MRI experiments. AVA performed fiber tractography. All authors reviewed and edited the manuscript.

## References

[1] Lichtman Jeff W., Denk Winfried. The big and the small: Challenges of imaging the brain’s circuits. Science. 2011;334(6056):618–623.

[2] Basser Peter J., Mattiello James, LeBihan Denis. Estimation of the Effective Self-Diffusion Tensor from the NMR Spin Echo. Journal of Magnetic Resonance, Series B. 1994;103(3):247–254.

[3] Basser P.J., Mattiello J., LeBihan D.. MR diffusion tensor spectroscopy and imaging. Biophysical Journal. 1994;66(1):259–267.

[4] Stejskal E. O., Tanner J. E.. Spin Diffusion Measurements: Spin Echoes in the Presence of a Time-Dependent Field Gradient. The Journal of Chemical Physics. 1965;42(1):288–292.

[5] Pierpaoli Carlo, Basser Peter J.. Toward a quantitative assessment of diffusion anisotropy. Magnetic Resonance in Medicine. 1996;36(6):893–906.

[6] Tuch David S., Reese Timothy G., Wiegell Mette R., Wedeen Van J.. Diffusion MRI of Complex Neural Architecture. Neuron. 2003;40(5):885–895.

[7] Özarslan Evren, Mareci Thomas H.. Generalized diffusion tensor imaging and analytical relationships between diffusion tensor imaging and high angular resolution diffusion imaging. Magnetic Resonance in Medicine. 2003;50(5):955–965.

[8] Cheng Yuan, Cory David G.. Multiple scattering by NMR. Journal of the American Chemical Society. 1999;121(34):7935–7936.

[9] Cory DG, Garroway Allen N., Miller Joel B.. Applications of spin transport as a probe of local geometry. In: Proceedings of the American Chemical Society:149–150; 1990.

[10] Callaghan P. T., Komlosh M. E.. Locally anisotropic motion in a macroscopically isotropic system: displacement correlations measured using double pulsed gradient spin-echo NMR. Magnetic Resonance in Chemistry. 2002;40(13):S15–S19.

[11] Komlosh M.E., Horkay F., Freidlin R.Z., Nevo U., Assaf Yaniv, Basser Peter J.. Detection of microscopic anisotropy in gray matter and in a novel tissue phantom using double Pulsed Gradient Spin Echo MR. Journal of Magnetic Resonance. 2007;189(1):38–45.

[12] Avram Alexandru V., Özarslan Evren, Sarlls Joelle E., Basser Peter J.. In vivo detection of microscopic anisotropy using quadruple pulsed-field gradient (qPFG) diffusion MRI on a clinical scanner. NeuroImage. 2013;64:229–239.

[13] Lawrenz Marco, Finsterbusch Jürgen. Detection of microscopic diffusion anisotropy in human cortical gray matter in vivo with double diffusion encoding. Magnetic Resonance in Medicine. 2019;81(2):1296–1306.

[14] Jespersen Sune Nørhøj, Lundell Henrik, Sønderby Casper Kaae, Dyrby Tim B.. Orientationally invariant metrics of apparent compartment eccentricity from double pulsed field gradient diffusion experiments. NMR in Biomedicine. 2013;26(12):1647–1662.

[15] Koch Martin A., Finsterbusch Jürgen. Compartment size estimation with double wave vector diffusion-weighted imaging. Magnetic Resonance in Medicine. 2008;60(1):90–101.

[16] Komlosh M.E., Lizak M.J., Horkay F., Freidlin R.Z., Basser Peter J.. Observation of microscopic diffusion anisotropy in the spinal cord using double-pulsed gradient spin echo MRI. Magnetic Resonance in Medicine. 2008;59(4):803–809.

[17] MacAulay N., Hamann S., Zeuthen T.. Water transport in the brain: Role of cotransporters. Neuroscience. 2004;129(4):1029–1042.

[18] Basser P.J., Pajevic S.. A normal distribution for tensor-valued random variables: applications to diffusion tensor MRI. IEEE Transactions on Medical Imaging. 2003;22(7):785–794.

[19] Jian Bing, Vemuri Baba C, Ozarslan Evren, Carney Paul R., Mareci Thomas H. A novel tensor distribution model for the diffusion-weighted MR signal.. NeuroImage. 2007;37(1):164–76.

[20] Westin Carl-Fredrik, Knutsson Hans, Pasternak Ofer, et al. Q-space trajectory imaging for multidimensional diffusion MRI of the human brain. NeuroImage. 2016;135:345–362.

[21] Reymbaut Alexis. Matrix moments of the diffusion tensor distribution and matrix-variate Gamma approximation. Journal of Magnetic Resonance Open. 2021;8-9:100016.

[22] Topgaard Daniel. Multidimensional diffusion MRI. Journal of Magnetic Resonance. 2017;275:98–113.

[23] Song Yiqiao, Ly Ina, Fan Qiuyun, et al. Measurement of Full Diffusion Tensor Distribution Using High-Gradient Diffusion MRI and Applications in Diffuse Gliomas. Frontiers in Physics. 2022;0:196.

[24] Magdoom Kulam Najmudeen, Pajevic Sinisa, Dario Gasbarra, Basser Peter J. A new framework for MR diffusion tensor distribution. Scientific Reports. 2021;11(1):2766.

[25] Avram Alexandru V., Sarlls Joelle E., Basser Peter J.. Measuring non-parametric distributions of intravoxel mean diffusivities using a clinical MRI scanner. NeuroImage. 2019;185:255–262.

[26] Wong Eric C., Cox Robert W., Song Allen W.. Optimized isotropic diffusion weighting. Magnetic Resonance in Medicine. 1995;34(2):139–143.

[27] Wedeen Van J., Rosene Douglas L., Wang Ruopeng, et al. The geometric structure of the brain fiber pathways. Science. 2012;335(6076):1628–1634.

[28] Avram Alexandru V, Sarlls Joelle E, Basser Peter J, Ozarslan Evren. Rotating Field Gradient (RFG) Diffusion MRI for Mapping 3D Orientation Distribution Functions (ODFs) in the Human Brain. In: Proc Intl Soc Mag Reson Med:6644–6644; 2014.

[29] Szczepankiewicz Filip, Westin Carl-Fredrik, Nilsson Markus. Maxwell-compensated design of asymmetric gradient waveforms for tensor-valued diffusion encoding. Magnetic Resonance in Medicine. 2019;82(4):1424–1437.

[30] Topgaard Daniel. Isotropic diffusion weighting in PGSE NMR: Numerical optimization of the q-MAS PGSE sequence. Microporous and Mesoporous Materials. 2013;178:60–63.

[31] Sjölund Jens, Szczepankiewicz Filip, Nilsson Markus, Topgaard Daniel, Westin Carl-Fredrik, Knutsson Hans. Constrained optimization of gradient waveforms for generalized diffusion encoding. Journal of Magnetic Resonance. 2015;261:157–168.

[32] Mitra Partha P.. Multiple wave-vector extensions of the NMR pulsed-field-gradient spin-echo diffusion measurement. Physical Review B. 1995;51(21):15074–15078.

[33] Zhou Xiaohong Joe, Tan Steve G., Bernstein Matt A.. Artifacts induced by concomitant magnetic field in fast spin-echo imaging. Magnetic Resonance in Medicine. 1998;40(4):582–591.

[34] Bernstein Matt A., Zhou X J, Polzin J a, et al. Concomitant gradient terms in phase contrast MR: analysis and correction.. Magnetic resonance in medicine. 1998;39(2):300–8.

[35] Baron C. A., Lebel R. M., Wilman A. H., Beaulieu C.. The effect of concomitant gradient fields on diffusion tensor imaging. Magnetic Resonance in Medicine. 2012;68(4):1190–1201.

[36] Veraart Jelle, Novikov Dmitry S., Christiaens Daan, Ades-aron Benjamin, Sijbers Jan, Fieremans Els. Denoising of diffusion MRI using random matrix theory. NeuroImage. 2016;142:394–406.

[37] Garyfallidis Eleftherios, Brett Matthew, Amirbekian Bagrat, et al. Dipy, a library for the analysis of diffusion MRI data. Frontiers in Neuroinformatics. 2014;8(FEB):8.

[38] Jenkinson M, Smith S. A global optimisation method for robust affine registration of brain images.. Medical image analysis. 2001;5(2):143–56.

[39] Jenkinson Mark, Bannister Peter, Brady Michael, Smith Stephen. Improved optimization for the robust and accurate linear registration and motion correction of brain images.. NeuroImage. 2002;17(2):825–41.

[40] Smith Stephen M., Jenkinson Mark, Woolrich Mark W., et al. Advances in functional and structural MR image analysis and implementation as FSL. NeuroImage. 2004;23(SUPPL. 1):S208–S219.

[41] Andersson Jesper L.R., Skare Stefan, Ashburner John. How to correct susceptibility distortions in spin-echo echo-planar images: application to diffusion tensor imaging.. NeuroImage. 2003;20(2):870–88.

[42] Freidlin Raisa Z., Özarslan Evren, Komlosh Michal E., et al. Parsimonious model selection for tissue segmentation and classification applications: A study using simulated and experimental DTI data. IEEE Transactions on Medical Imaging. 2007;26(11):1576–1584.

[43] Basser Peter J., Pajevic Sinisa. Spectral decomposition of a 4th-order covariance tensor: Applications to diffusion tensor MRI. Signal Processing. 2007;87(2):220–236.

[44] Ennis Daniel B., Kindlmann Gordon. Orthogonal tensor invariants and the analysis of diffusion tensor magnetic resonance images. Magnetic Resonance in Medicine. 2006;55(1):136–146.

[45] Descoteaux Maxime, Deriche Rachid, Knösche Thomas R., Anwander Alfred. Deterministic and probabilistic tractography based on complex fibre orientation distributions. IEEE Transactions on Medical Imaging. 2009;28(2):269–286.

[46] Tournier J-Donald, Calamante Fernando, Connelly Alan. MRtrix: Diffusion tractography in crossing fiber regions. International Journal of Imaging Systems and Technology. 2012;22(1):53–66.

[47] Jensen Jens H., Helpern Joseph A., Ramani Anita, Lu Hanzhang, Kaczynski Kyle. Diffusional kurtosis imaging: The quantification of non-gaussian water diffusion by means of magnetic resonance imaging. Magnetic Resonance in Medicine. 2005;53(6):1432–1440.

[48] Fieremans Els, Burcaw Lauren M., Lee Hong Hsi, Lemberskiy Gregory, Veraart Jelle, Novikov Dmitry S.. In vivo observation and biophysical interpretation of time-dependent diffusion in human white matter. NeuroImage. 2016;129:414–427.

[49] Tuch David S, Reese Timothy G, Wiegell Mette R, Makris Nikos, Belliveau John W., Wedeen Van J. High angular resolution diffusion imaging reveals intravoxel white matter fiber heterogeneity.. Magnetic resonance in medicine. 2002;48(4):577–82.

[50] Assaf Yaniv, Basser Peter J.. Composite hindered and restricted model of diffusion (CHARMED) MR imaging of the human brain. NeuroImage. 2005;27(1):48–58.

[51] Assaf Yaniv, Blumenfeld-Katzir Tamar, Yovel Yossi, Basser Peter J.. Axcaliber: A method for measuring axon diameter distribution from diffusion MRI. Magnetic Resonance in Medicine. 2008;59(6):1347–1354.

[52] Basser Peter J. Fiber-Tractography via Diffusion Tensor MRI (DT-MRI). In: Proceedings of the 6th Annual Meeting ISMRM:1226; 1998; Sydney, Australia.

[53] Mori Susumu, Crain Barbara J., Zijl Peter C.M. 3D Brain Fiber Reconstruction from Diffusion MRI. NeuroImage. 1998;7(4):S710.

[54] Tournier J. Donald, Yeh Chun Hung, Calamante Fernando, Cho Kuan Hung, Connelly Alan, Lin Ching Po. Resolving crossing fibres using constrained spherical deconvolution: Validation using diffusion-weighted imaging phantom data. NeuroImage. 2008;42(2):617–625.

[55] Jbabdi S., Woolrich M.W., Andersson J.L.R., Behrens T.E.J.. A Bayesian framework for global tractography. NeuroImage. 2007;37(1):116–129.

[56] Smith Robert E., Tournier Jacques Donald, Calamante Fernando, Connelly Alan. SIFT: Spherical-deconvolution informed filtering of tractograms. NeuroImage. 2013;67:298–312.

[57] Tournier J. Donald, Calamante Fernando, Gadian David G., Connelly Alan. Diffusion-weighted magnetic resonance imaging fibre tracking using a front evolution algorithm. NeuroImage. 2003;20(1):276–288.

[58] Jeurissen Ben, Szczepankiewicz Filip. Multi-tissue spherical deconvolution of tensor-valued diffusion MRI. NeuroImage. 2021;245:118717.

[59] Dell’Acqua Flavio, Tournier J.-Donald. Modelling white matter with spherical deconvolution: How and why?. NMR in Biomedicine. 2019;32(4):e3945.

[60] Shapson-Coe Alexander, Januszewski Michal, Berger Daniel R., et al. A connectomic study of a petascale fragment of human cerebral cortex. bioRxiv. 2021;:2021.05.29.446289.

[61] Abdollahzadeh Ali, Belevich Ilya, Jokitalo Eija, Sierra Alejandra, Tohka Jussi. DeepACSON automated segmentation of white matter in 3D electron microscopy. Communications Biology 2021 4:1. 2021;4(1):1–14.

[62] Pasternak Ofer, Sochen Nir, Gur Yaniv, Intrator Nathan, Assaf Yaniv. Free water elimination and mapping from diffusion MRI. Magnetic Resonance in Medicine. 2009;62(3):717–730.

[63] Liewald Daniel, Miller Robert, Logothetis Nikos, Wagner Hans Joachim, Schüz Almut. Distribution of axon diameters in cortical white matter: an electron-microscopic study on three human brains and a macaque. Biological Cybernetics. 2014;108(5):541–557.

[64] Thorne Robert G., Nicholson Charles. In vivo diffusion analysis with quantum dots and dextrans predicts the width of brain extracellular space. Proceedings of the National Academy of Sciences of the United States of America. 2006;103(14):5567–5572.

[65] Malmborg Carin, Sjöbeck Martin, Brockstedt Sara, Englund Elisabeth, Söderman Olle, Topgaard Daniel. Mapping the intracellular fraction of water by varying the gradient pulse length in q-space diffusion MRI. Journal of Magnetic Resonance. 2006;180:280–285.

[66] McNab Jennifer A., Edlow Brian L., Witzel Thomas, et al. The Human Connectome Project and beyond: Initial applications of 300mT/m gradients. NeuroImage. 2013;80:234–245.

[67] Foo Thomas K. F., Tan Ek Tsoon, Vermilyea Mark E., et al. Highly efficient head-only magnetic field insert gradient coil for achieving simultaneous high gradient amplitude and slew rate at 3.0T (MAGNUS) for brain microstructure imaging. Magnetic Resonance in Medicine. 2020;83(6):2356–2369.

[68] Huang Susie Y., Witzel Thomas, Keil Boris, et al. Connectome 2.0: Developing the next-generation ultra-high gradient strength human MRI scanner for bridging studies of the micro-, meso- and macro-connectome. NeuroImage. 2021;:118530.

[69] Quirk James D., Bretthorst G. Larry, Duong Timothy Q., et al. Equilibrium water exchange between the intra- and extracellular spaces of mammalian brain. Magnetic Resonance in Medicine. 2003;50(3):493–499.

[70] Bai Ruiliang, Springer Charles S., Plenz Dietmar, Basser Peter J.. Fast, Na + /K + pump driven, steady-state transcytolemmal water exchange in neuronal tissue: A study of rat brain cortical cultures. Magnetic Resonance in Medicine. 2018;79(6):3207–3217.

[71] Williamson Nathan H., Ravin Rea, Benjamini Dan, et al. Magnetic resonance measurements of cellular and sub-cellular membrane structures in live and fixed neural tissue. eLife. 2019;8.

[72] Chen Liyong, Beckett Alexander, Verma Ajay, Feinberg David A. Dynamics of respiratory and cardiac CSF motion revealed with real-time simultaneous multi-slice EPI velocity phase contrast imaging. NeuroImage. 2015;122:281–287.

[73] Avram Alexandru V., Sarlls Joelle E., Basser Peter J.. Whole-Brain Imaging of Subvoxel T1-Diffusion Correlation Spectra in Human Subjects. Frontiers in Neuroscience. 2021;15:682.

[74] Yang Grant, Tian Qiyuan, Leuze Christoph, Wintermark Max, McNab Jennifer A.. Double diffusion encoding MRI for the clinic. Magnetic Resonance in Medicine. 2018;80(2):507–520.

[75] Axer Hubertus, Keyserlingk Diedrich Graf V.. Mapping of fiber orientation in human internal capsule by means of polarized light and confocal scanning laser microscopy. Journal of Neuroscience Methods. 2000;94(2):165–175.

[76] Tax Chantal M.W., Szczepankiewicz Filip, Nilsson Markus, Jones Derek K.. The dot-compartment revealed? Diffusion MRI with ultra-strong gradients and spherical tensor encoding in the living human brain. NeuroImage. 2020;210:116534.

