## Supplementary Material for "A Novel Framework for *In-vivo* Diffusion Tensor Distribution MRI of the Human Brain"

### 1 Rank of b-tensor generated by iPFG

The effective gradient vector, $\mathbf{G}(t)$, in diffusion module with two interfused pulsed field gradients (iPFG), $\mathbf{g}_1(t)$ and $\mathbf{g}_2(t)$, accounting for phase inversion by the 180-degree RF pulse is given by,

$$\mathbf{G}(t) = \begin{cases} -\mathbf{g}_1(t) & 0 \leq t \leq \delta \\ -\mathbf{g}_2(t-\delta) & \delta \leq t \leq 2\delta \\ 0 & 2\delta \leq t \leq \Delta \\ \mathbf{g}_1(t-\Delta) & \Delta \leq t \leq \Delta+\delta \\ \mathbf{g}_2(t-\Delta-\delta) & \Delta+\delta \leq t \leq \Delta+2\delta \end{cases} \quad (1)$$

A component of the b-tensor generated by the above gradient is given by,

$$\mathsf{b}_{ij} = \int_0^{\Delta+2\delta} \left(\gamma \int_0^t G_i(t')dt'\right) \left(\gamma \int_0^t G_i(t'')dt''\right) dt \quad (2)$$

where $G_i$ is the $i^{th}$-component of the gradient vector, $\mathbf{G}$. Plugging in the effective gradient Equation (1) into the above relation, the expression for $\mathsf{b}_{ij}$ is given by,

$$\mathsf{b}_{ij} = \frac{1}{3}\gamma^2\delta^2 \left(-(g_{2i}(3g_{1j}+g_{2j}) + g_{1i}(g_{1j}+3g_{2j}))\delta + 3(g_{1i}+g_{2i})(g_{1j}+g_{2j})\Delta\right) \quad (3)$$

The rank-2 b-tensor generated by such a interfused gradient pattern is demonstrated by plugging in $\mathbf{g}_1 = [1, 0, 0]$ and $\mathbf{g}_2 = [0, 1, 0]$ into the above expression resulting in the following b-tensor,

$$\mathsf{b} = \frac{\gamma^2 \delta^2}{3} \begin{bmatrix} 3\Delta - \delta & 3(\Delta - \delta) & 0 \\ 3(\Delta - \delta) & 3\Delta - \delta & 0 \\ 0 & 0 & 0 \end{bmatrix} \tag{4}$$

The above matrix has two non-zero eigenvalues $\left(\text{i.e.,} \frac{2\gamma^2\delta^3}{3}, \frac{2}{3}\gamma^2\delta^2(3\Delta - 2\delta)\right)$ indicating its rank is 2.

### 2 $q$-space formalism using iPFG for restricted diffusion

The echo attenuation for our pulse sequence is given by,

$$E(\mathbf{q}_1, \mathbf{q}_2) = \int\int\int\int \rho(\mathbf{r}) \exp\left(-i\mathbf{q}_1 \cdot \mathbf{r}\right) P(\mathbf{r}|\mathbf{r}', \delta) \exp\left(-i\mathbf{q}_2 \cdot \mathbf{r}'\right) P(\mathbf{r}'|\mathbf{r}'', \Delta) \exp\left(i\mathbf{q}_1 \cdot \mathbf{r}''\right)$$
$$P(\mathbf{r}''|\mathbf{r}''', \delta) \exp\left(i\mathbf{q}_2 \cdot \mathbf{r}'''\right) d\mathbf{r}d\mathbf{r}'d\mathbf{r}''d\mathbf{r}''' \tag{5}$$

where $\mathbf{q}_1, \mathbf{q}_2$ are the time integral under the two independent diffusion gradient pulses, $\mathbf{r}$ is the spin position, $\rho(\mathbf{r})$ is the initial spin density at $\mathbf{r}$ and $P(\mathbf{r}_1|\mathbf{r}_2, dt)$ is the propagator which is the conditional probability for the spins to move from $\mathbf{r}_1$ to $\mathbf{r}_2$ in time interval, $dt$.

Assuming restricted diffusion within the pore of size, $a$, and spin diffusivity, $D$, simplified expressions can be obtained for the above integral in the two time regimes which look very similar to those obtained from traditional dPFG sequence.

#### 2.1 At $\delta \gg a^2/D$

In the limit, the mixing (i.e., $\delta$) time of the experiment is much greater than the time required to explore the pore, the propagator reduces to spin density of the pore resulting in the following expression for the echo attenuation,

$$E(\mathbf{q}_1, \mathbf{q}_2) = \int\int\int\int \rho(\mathbf{r}) \exp\left(-i\mathbf{q}_1 \cdot \mathbf{r}\right) \rho(r') \exp\left(-i\mathbf{q}_2 \cdot \mathbf{r}'\right) \rho(r'') \exp\left(i\mathbf{q}_1 \cdot \mathbf{r}''\right) \rho(r''')$$
$$\exp\left(i\mathbf{q}_2 \cdot \mathbf{r}'''\right) d\mathbf{r}d\mathbf{r}'d\mathbf{r}''d\mathbf{r}''' \tag{6}$$

Rearranging the integrals,

$$E(\mathbf{q}_1, \mathbf{q}_2) = \left|\int \rho(\mathbf{r}) \exp\left(i\mathbf{q}_1 \cdot \mathbf{r}\right) d\mathbf{r}\right|^2 \left|\int \rho(\mathbf{r}) \exp\left(i\mathbf{q}_2 \cdot \mathbf{r}\right) d\mathbf{r}\right|^2 \tag{7}$$
$$= |\tilde{\rho}(\mathbf{q}_1)|^2 |\tilde{\rho}(\mathbf{q}_2)|^2 \tag{8}$$

where $\tilde{\rho}(\mathbf{q})$ is the pore spectral density.

### 2.2 At $\Delta \ll a^2/D$

In the limit, the diffusion (i.e., $\Delta$) time of the experiment is much smaller than the time required to explore the pore, the propagator reduces to a delta function resulting in the following expression for the echo attenuation,

$$E(\mathbf{q}_1, \mathbf{q}_2) = \int\int\int\int \rho(\mathbf{r}) \exp\left(-i\mathbf{q}_1 \cdot \mathbf{r}\right) \delta(r - r') \exp\left(-i\mathbf{q}_2 \cdot \mathbf{r}'\right) \delta(r' - r'') \exp\left(i\mathbf{q}_1 \cdot \mathbf{r}''\right) \delta(r'' - r''') \exp\left(i\mathbf{q}_2 \cdot \mathbf{r}'''\right) d\mathbf{r} d\mathbf{r}' d\mathbf{r}'' d\mathbf{r}''' \quad (9)$$

Rearranging the integrals,

$$E(\mathbf{q}_1, \mathbf{q}_2) = \left| \int \rho(\mathbf{r}) \exp\left(i(\mathbf{q}_1 + \mathbf{q}_2) \cdot \mathbf{r}\right) d\mathbf{r} \right|^2 \quad (10)$$

$$= |\tilde{\rho}(\mathbf{q}_1 + \mathbf{q}_2)|^2 \quad (11)$$

A equivalent relation between echo attenuation and polar angle between the gradients obtained in traditional dPFG sequences is derived as follows. Expanding the pore spectral density function using Taylor series and assuming inversion symmetry,

$$\tilde{\rho}(\mathbf{q}) \approx 1 - \frac{1}{2} \int \rho(\mathbf{r}) (\mathbf{q} \cdot \mathbf{r})^2 d\mathbf{r} + \mathcal{O}(q^4) \quad (12)$$

Applying the above relation to the derived echo attenuation expression and assuming $\mathbf{q}_1, \mathbf{q}_2$ are of equal amplitude, $q$, and makes a polar angle, $\theta_1$ and $\theta_2$ respectively with respect to $\mathbf{r}$,

$$\tilde{\rho}(\mathbf{q}_1 + \mathbf{q}_2) \approx 1 - \frac{1}{2} \int \rho(\mathbf{r}) ((\mathbf{q}_1 + \mathbf{q}_2) \cdot \mathbf{r})^2 d\mathbf{r} \quad (13)$$

$$= 1 - \frac{q^2}{2} \int r^2 \rho(\mathbf{r}) (\cos\theta_1 + \cos\theta_2)^2 d\mathbf{r} \quad (14)$$

$$= 1 - 2q^2 \cos^2\left(\frac{\theta_1 - \theta_2}{2}\right) \int r^2 \rho(\mathbf{r}) \cos^2\left(\frac{\theta_1 + \theta_2}{2}\right) d\mathbf{r} \quad (15)$$

$$= 1 - q^2 (1 + \cos(\Theta)) \int r^2 \rho(\mathbf{r}) \cos^2\theta_{12} d\mathbf{r} \quad (16)$$

Where $\Theta = \theta_1 - \theta_2$ is the polar angle between the two $q$-vectors, and $\theta_{12} = \frac{\theta_1 + \theta_2}{2}$. Substituting this expression in the echo attenuation above,

$$E(\mathbf{q}_1, \mathbf{q}_2) = |\tilde{\rho}(\mathbf{q}_1 + \mathbf{q}_2)|^2 = 1 - 2q^2 (1 + \cos(\Theta)) \int r^2 \rho(\mathbf{r}) \cos^2\theta_{12} d\mathbf{r} + \mathcal{O}(q^4) \quad (17)$$
